# Private Genomes and Public SNPs: Homomorphic encryption of genotypes and phenotypes for shared quantitative genetics

**DOI:** 10.1101/2020.04.02.021865

**Authors:** Richard Mott, Christian Fischer, Pjotr Prins, Robert William Davies

**Affiliations:** Genetics Institute, University College London, Gower St London WC1E 6BT; Genetics, Genomics and Informatics, University of Tennessee Health Science Center, 71 S Manassas St, Memphis TN, USA; Department of Statistics, University of Oxford, 29 St Giles’, Oxford OX1 3LB, UK

## Abstract

Sharing human genotype and phenotype data presents a challenge because of privacy concerns, but is essential in order to discover otherwise inaccessible genetic associations. Here we present a method of homomorphic encryption that obscures individuals’ genotypes and phenotypes and is suited to quantitative genetic association analysis. Encrypted ciphertext and unencrypted plaintext are interchangeable from an analytical perspective. This allows one to store ciphertext on public web services and share data across multiple studies, while maintaining privacy. The encryption method uses as its key a high-dimensional random linear orthogonal transformation that leaves the likelihood of quantitative trait data unchanged under a linear model with normally distributed errors. It also preserves linkage disequilibrium between genetic variants and associations between variants and phenotypes. It scrambles relationships between individuals: encrypted genotype dosages closely resemble Gaussian deviates, and in fact can be replaced by quantiles from a Gaussian with only negligible effects on accuracy. Standard likelihood-based inferences are unaffected by orthogonal encryption. These include the use of mixed linear models to control for unequal relatedness between individuals, the estimation of heritability, and the inclusion of covariates when testing for association. Orthogonal transformations can also be applied in a modular fashion that permits multi-party federated mega-analyses. Under this scheme any number of parties first agree to share a common set of genotype sites and covariates prior to encryption. Each party then privately encrypts and shares their own ciphertext, and analyses the other parties’ ciphertexts. In the absence of private variants, or knowledge of the key, we show that it is infeasible to decrypt ciphertext using existing brute-force or noise reduction attacks. Therefore, we present the method as a challenge to the community to determine its security.

## Introduction

With the growth of clinical genome sequencing, the numbers of individual human genomes available for analysis is expected to increase dramatically. To make the most of this resource we need to be able to share and analyse genetic and phenotypic data securely, and the conflicting demands of individual privacy and medical research have led to a spectrum of ways of sharing human genotype and phenotype data(Azencott 2018).

In a small minority of studies, anonymised data (that is, where the names of individuals have been replaced by anonymous identifiers) are freely available for users to download and analyse. More usually - as for the UK BioBank and UK 10k project, and studies deposited in NCBI dbGAP and the EBI GPA - anonymised data are distributed only to researchers approved for access, and whose institutions demonstrate that their computer systems are secure, and where they agree not to redistribute the data. The host data archive then prepares datasets, encrypted with keys that may be specific to each data request, for transfer over a public network. After downloading the encrypted files within the firewall of the researcher’s computer system, they are decrypted into plain text. The advantage of this approach is that the researcher then has complete access to the anonymised genotypes and phenotypes, with only the identities of the samples being redacted; there is then no technical limitation as to the genetic analysis that can be performed. However, this carries certain risks because a data breach cannot be ruled out, and even if the data are anonymised, comparing anonymous genotypes with those of genotyped relatives might still reveal genetic relationships(Hansson *et al.* 2016). In the clinical field, methods such as the random time-shifting of anonymised patient records(Hripcsak *et al.* 2016) offer some protection whilst not being cryptographically secure.

At the other extreme, datasets are not distributed, but researchers may negotiate access to analyse the data on the host’s computer system (as in the UK 100,000 genomes project), or the host may agree to perform an analysis on behalf of an external user. No direct access to the raw data is granted, but analyses are shared. In still other cases, only the summary statistics of Genome Wide Association Studies (GWAS) are distributed, typically comprising the regression coefficients and p-values of the genetic variants tested for association with the phenotype, for a federated *meta-analysis.* Such analyses combine sets of summary statistics from different GWAS, where participating laboratories have collected phenotypes and genotypes for different sets of subjects imputed at the same SNPs, and wish to test association across all studies(Pasaniuc and Price 2017).

Another approach that is gaining traction is to encrypt genotypes and phenotypes in such a way that it is still possible to perform relevant computations on the data - possibly on a remote or cloud computer - without decrypting them, i.e., one can ‘throw away the key’. Homomorphic encryption (HE) are cryptographic systems that allow computations to be performed on encrypted data (the ciphertext) without decrypting, it and which yield the same answers as when the analogous computations are performed on the original data (the plaintext). It is an active area of research in computer science because it could make cloud computing much more secure, for both genetic and other applications. With HE, it is possible to build systems that store and process encrypted data, such that the data always stays encrypted both in transit and at rest. Should a cloud service be compromised, any stolen ciphertext would be valueless.

We define Homomorphic Encryption for Genotypes and Phenotypes (HEGP) to mean a transformation of the data that preserves the structure necessary for analysis whilst obscuring the individuals’ identities, phenotypes and genotypes. Only the encrypted data is moved and shared between systems. HEGP is attractive because it enables testing genetic association across multiple data sets, in a federated *mega-analysis* based on the genotypes instead of a less powerful meta-analysis based on the summary statistics.

In statistical genetics, a number of approaches to HEGP have been proposed. In(Jagadeesh *et al.* 2017) Yao’s protocol is used to identify rare Mendelian-type mutations shared between affected individuals. In(Cho *et al.* 2018) secure multi-party communication is used to perform GWAS using principal components to control for population structure. (Bonte *et al.* 2018; Tkachenko *et al.* 2018; Sim *et al.* 2019) describe cryptographically secure protocols for computing P-values for case-control studies using contingency table chi-squared tests. All these methods are thought to be cryptographically secure, but they limit the types of computation and data exploration possible. In particular, they cannot control for population structure using a mixed linear model, which is the current gold standard for quantitative trait analysis. In addition, they tend to be slower than analyses of un-encrypted data.

Here we consider whether linear transformations of genotypes and phenotypes can be used as keys for homomorphic encryption. The first class of transformations we investigate are random orthogonal transformations. These leave invariant essential parts of the linear mixed model framework for complex trait analysis commonly used in quantitative genetics, preserving genotype correlations between Single Nucleotide Polymorphisms (SNPs) whilst obscuring those between individuals. They share the same likelihood functions as un-encrypted data. Any standard mixed-model type of analysis (including estimating heritability) will produce the same output as with unencrypted data. We ask if an orthogonal key can be generated in a sufficiently random manner to make the data unrecognizable, and show that keys sampled from the Steiffel manifold have this property: however, not all orthogonal matrices make suitable keys. Once encryption has taken place, we show computations are essentially identical to those using unencrypted data. They also can be extended to perform federated mega-analyses in a natural way. Their major drawbacks are that they are unsuitable for logistic regression, and that the method is not provably secure. In particular, individuals with private variants are not securely encrypted by orthogonal transformation. However, for variants present in multiple individuals we present arguments that suggest it would be very challenging to find the key and hence decrypt the data.

The second type of linear transformation we consider is based on the mixed-model transformation. We show that this is likely to be more secure than orthogonal transformation but is more limited in its applications.

## METHODS AND RESULTS

### Conceptual Overview

The conflict between respecting individuals’ privacy and establishing allelic effects is sketched in **Figure 1A**. We have a vector of phenotypes ***y*** and a matrix of genotypes, ***G***. Each row of the matrix corresponds to genotypes for a given individual, and each column to a given SNP. The phenotype and each genotype vector (column of ***G***) is standardised to have mean 0 and variance 1. The genotypes are dosages proportional to the estimated number of alternative alleles; a typical trimodal distribution of dosages is also shown in **Figure 1C**. We want to preserve the privacy of the individuals (rows) but make public certain information about the effects of the SNPs (columns) in relation to each other and to the phenotype.

**Figure 1.**
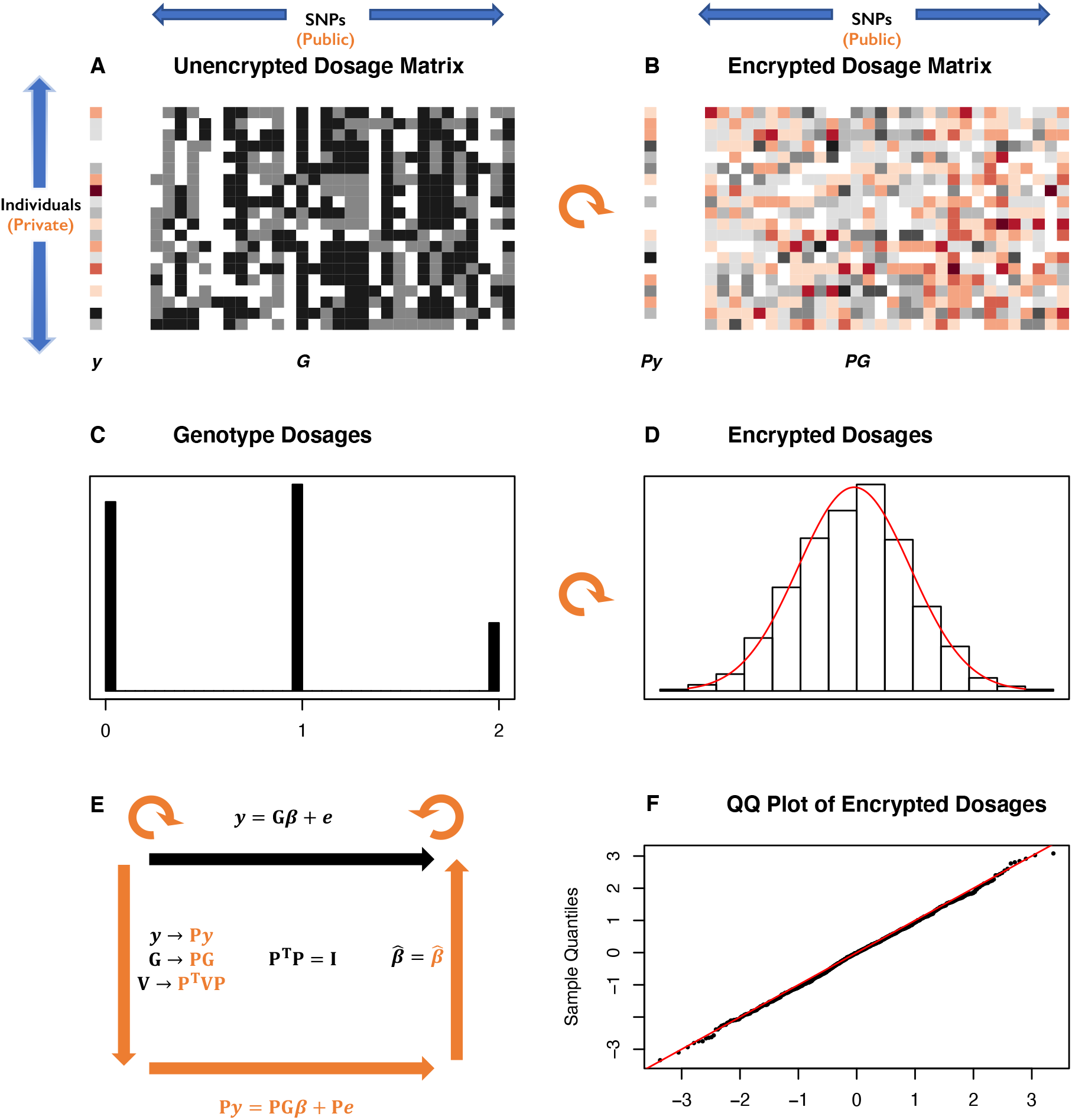
Privacy in relation to genetic association testing. **A:** A phenotype vector **y** (left) and genotype matrix **G** (right) are represented as colours and shades of grey. Each row of the matrix is one individual and each column one SNP. Genotypes are encoded as imputed dosages clustered at the values 0,1,2 giving the numbers of minor alleles. A typical distribution of dosages for one SNP is shown to the right. The aim is to hide information about rows but make public the relationships between the columns and the phenotype. **B:** the same data after multiplication by an orthogonal matrix P (a rotation represented by the curved orange arrow). The genotype dosages are now represented by a continuum of real numbers. **C:** The distribution of dosages for a particular SNP (column of G), clustered around 0,1,2. **D:** The distribution of the same dosages after orthogonal transformation by **P** (black histogram) with the Normal distribution with same mean and variance superimposed in red. F: the Normal qq-plot for the data in D, showing the transformed dosages are very close to a Normal distribution. **E:** A cartoon of the HEGP scheme. The top black arrow and equation shows the linear mixed model relating the phenotype **y** to genotype **G** with regression coefficients **β** representing the allelic effects. The variance matrix for the residuals is **V**. After multiplication by orthogonal matrix P, the data **y**, **G** and **V** and the mixed linear model are transformed as shown in orange. The likelihood and regression estimates 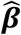 are preserved.

Conceptually, it is helpful to recall that the standardised genotype dosages for a given SNP across *n* subjects (a column in **Figure 1A**) can be thought of geometrically as a unit vector in *n*-dimensional space lying on the unit-dimensional hypersphere, and the vector of phenotypes as another point on the same hypersphere. We measure the association between phenotype and SNP from the angle *θ* between their *n*-dimensional vectors. Their Pearson correlation coefficient (an invertible transformation of the t-statistic used to determine significance of a linear regression of phenotype on genotype dosage) is equal to their dot-product, i.e. *cosθ*. Similarly, linkage disequilibrium *R*^2^ between any pair of SNPs is the square of the cosine of the angle between the SNPs. It is intuitively obvious that any orthogonal transformation – a rotation or reflection of the space - will leave all the angles between unit vectors unchanged (**Figure 2**). Thus all the associations between phenotype and genotypes, and correlations within genotypes, are preserved by orthogonal transformations. **Figure 1B** shows the phenotypes and genotypes after orthogonal transformation. Even though the original distribution of the genotypes dosages is trimodal (**Figure 1C**) the transformed genotypes resemble a sample from a Normal distribution (**Figure 1D,F**).

**Figure 2.**
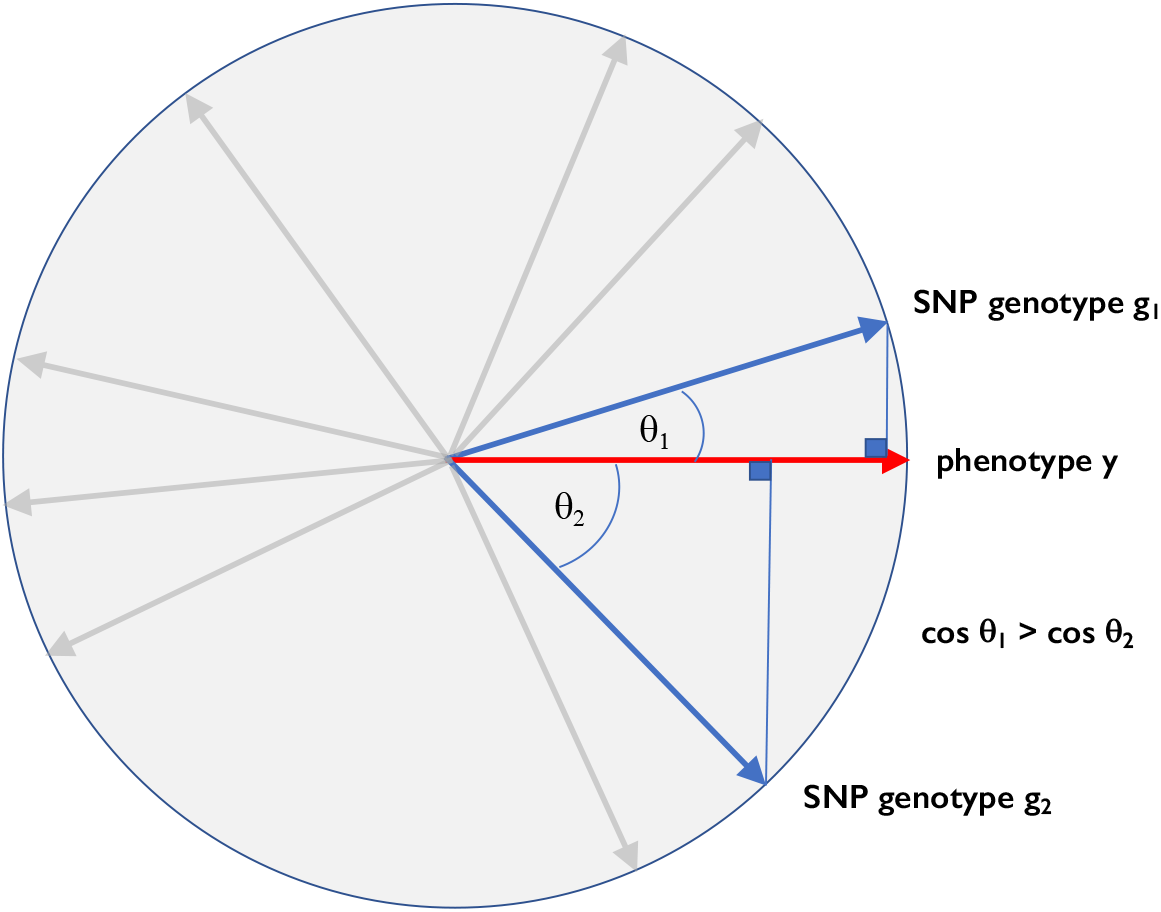
Geometric interpretation of genetic association. Phenotypes and genotypes are represented as vectors in a high-dimensional space. The cosines of the angles between the phenotype vector y and various SNPs equal the corresponding Pearson correlations, which are closely related to the t-statistics for testing association. In the example, SNP 1 has a smaller angle with the phenotype, than SNP 2, and hence a stronger genetic association.

It follows that, if the encryption key is an *n* × *n* orthogonal matrix ***P*** of floating point values such that ***PP***^***T***^ = ***I*** (where ***P***^***T***^ is the transpose of ***P***), then multiplication of the key with the genotype/phenotype matrix acts like a rotation (or reflection). In this way each SNP column is rotated by multiplication by the key, and as discussed below, if the key is sampled randomly, then the elements of each column vector of the resulting encrypted genotype/phenotype matrix are approximately normally distributed (**Figure 1D,F**). We next show that these transformations preserve key components of the linear mixed model relating the phenotype to the genotypes (**Figure 1E**)

### Statistical Preliminaries

#### Mixed model GWAS

In order to make this geometric intuition rigorous, we first review the core standard computations required for a mixed-model GWAS. Suppose we have a *n* subjects and *m* SNPs, a quantitative phenotype vector ***y*** of length *n*, a *n* × *p* covariate matrix ***X*** (containing information about e.g. sex, age, environmental covariates and principal components for controlling population structure) and a *n* × *m* genotype dosage matrix ***G*** in which the entries typically take the values 0,1,2, such that *G*_*ij*_ is the number of alternate alleles for the genotype of subject *i* at SNP *j* (***G*** can also represent imputed dosages without any change to the argument). It is necessary to standardise the genotype matrix into the matrix ***H*** such that

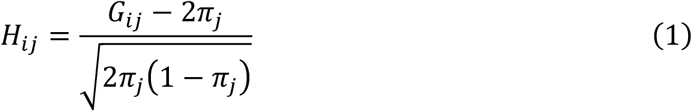

where *π*_*j*_ is the minor allele frequency of the SNP *j*. [Alternatively, each vector of dosages can be standardised empirically by subtracting its sample mean and dividing by its empirical standard deviation.] The phenotype vector ***y*** and each column of ***X*** must also be standardised to have mean 0 and variance 1.

The additive genetic relationship matrix

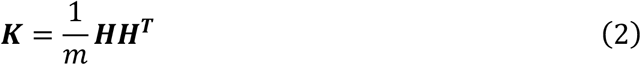

is used to model the variance-covariance structure of the phenotype as

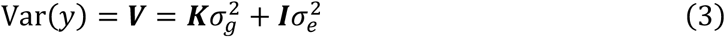

where 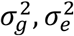 are the genetic and environmental variance components and

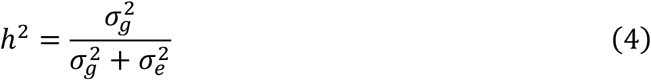

is the additive heritability. These variance components are typically estimated by restricted maximum-likelihood(Yang *et al.* 2011). The linear model to test the significance of the SNP *j* is

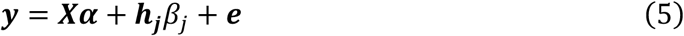

where ***α*** is a vector of fixed effects, ***h***_***j***_ is the *j*th column of ***H***,*β*_*j*_ is the regression coefficient for SNP *j* and ***e*** is the residual, with variance matrix ***V***.

The mixed model transformation

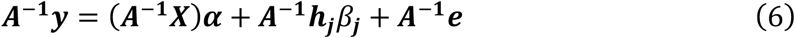

converts the mixed model into an Ordinary Least Squares problem in which the variance matrix is the identity, i.e. Var(***A***^**−1**^***V***) = ***I***. Here ***A*** is the matrix square root of ***V***, i.e. ***A***^2^ = ***V***, which can be computed efficiently by eigen-decomposition of ***K***, alongside the estimation of the variance components 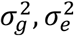 (Kang *et al.* 2008).

The genetic relationship between individuals *i*, *k* is summarised as *K*_*ik*_ and the relationship (Pearson correlation coefficient) between SNPs *j*, *l* as the element *L*_*jl*_ in the matrix

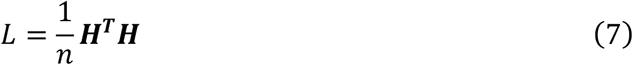

### Orthogonal Transformations

We wish to find an encoding of the genotypes, covariates and phenotype such that their plaintexts are obscured, but such that we can compute all the above quantities and test association between genotypes and phenotypes using the same mixed model.

Consider the eigen decomposition of the variance matrix ***V*** = ***E***^***T***^***ΛE*** where ***E*** is an orthonormal matrix of eigenvectors and ***Λ*** the diagonal matrix of eigenvalues. These quantities are determined (up to permutation and rotation) by the matrix ***V***. The (symmetric) matrix square root used in the mixed-model transformation is defined as

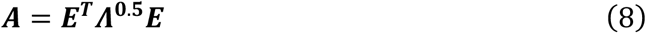

where ***Λ***^**0.5**^ is the diagonal matrix whose entries are the square roots of the eigenvalues.

Suppose ***P*** is *any* orthogonal *n* × *n* matrix, i.e. so that ***P***^**−1**^ = ***P***^***T***^. Then consider working with the transformed genotype matrix ***F*** = ***PH***, phenotype vector ***z*** = ***Py*** and covariate matrix ***W*** = ***PX*** in place of the plaintext. Such a transformation corresponds to finding a new coordinate system, so the rows (“subjects”) in the transformed space no longer correspond to individuals.

First note that

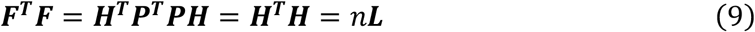

so the *m* × *m* SNP-relationship matrix ***L*** is preserved, while the *n* × *n* additive genetic relationship matrix, or GRM,

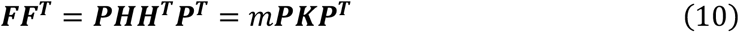

is transformed. In other words, linkage disequilibrium (as measured by Pearson correlation) between SNPs is unaltered, but since the original subjects are transformed, inter-subject correlations are destroyed. In fact, since after orthogonal transformation each “subject” is a weighted combination of the originals, it is not even meaningful to even describe them as subjects. Nonetheless,

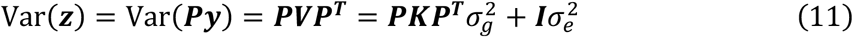

and hence the transformed phenotype has the same variance components 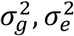 and heritability *h*^2^, even though the genetic relationship matrix is transformed. Define ***B*** = ***AP***. as the ***z***- analogue of the ***y*** mixed-model transformation. That is,

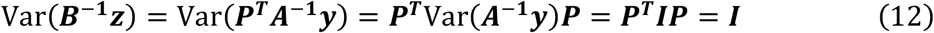

and hence the ciphertext “rotated mixed model”

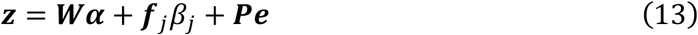

which is expressed entirely using the transformed quantities *D*(***P***) = {***z***, ***W***, ***F***} is equivalent to the original plaintext model and can be converted to ordinary least squares by multiplication by ***B***^**−1**^. Furthermore, the log-likelihood for the data (provided the errors are Normally distributed) is invariant after orthogonal transformation. That is, using standard change-of-variable rules for ***y*** = ***P***^***T***^***z*** for the multivariate normal distribution, and recalling that the determinant of an orthogonal matrix |***P***| = ±1, then the plaintext log likelihood for ***y***:

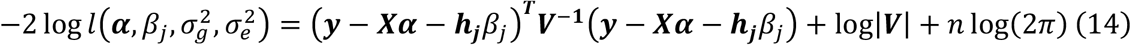

is identical to the log likelihood for ciphertext ***z*** when evaluated at the same parameters:

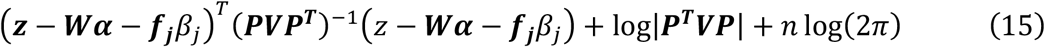

Hence all inferences about the parameters based on the likelihood are unaffected by the transformation. In particular they yield identical maximum likelihood parameter estimates and p-values for likelihood-based tests of significance. Furthermore, any analyses based on LD between SNPs are unaffected by the transformation. It is also possible to compute GRMs corresponding to subsets of SNPs (e.g. per-chromosome) from the transformed genotypes.

### Generalisations

Here we sketch various generalisations to the orthogonal encryption scheme.

(i) Analyses that are unaffected by orthogonal transformation include the estimation of parameters by **ridge regression** or by **Henderson’s mixed model equations**. The proof for ridge regression follows from the observation that the ridge estimator

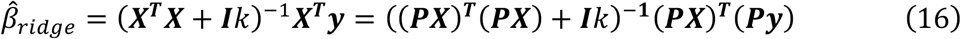

for any orthogonal matrix ***P*** and ridge scale parameter *k*. The proof for Henderson’s equations follows in a similar way, as under orthogonal transformation any data matrix transforms as ***X*** → ***PX*** and any variance matrix as ***V*** → ***PVP***^***T***^ (since Henderson’s model is a special case of a mixed model it also follows from Equation 6). Consequently, genomic prediction from estimated fixed effects (BLUE) and predicted random effects (BLUP) is also unaffected, provided of course we have access to some unencrypted genotypes with which to make predictions.

(ii) **Dominance effects** might be incorporated in the following way. The additive genotype dosage matrix ***G*** can be augmented in the usual way by a matrix ***T*** defined as

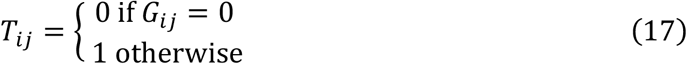

representing a dominance effect. Then any combination of additive and dominance effects can be modelled as a linear combination of ***G***, ***T***, so that Equation (5) that models the effect of SNP *j* becomes

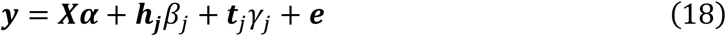

where ***t***_*j*_ is the jth column of T and *γ*_*j*_ is the dominance effect

Multiplying by the orthogonal matrix ***P*** produces

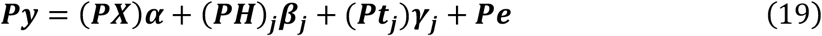

The rest of the development is similar to the purely additive case. Investigators would need to share both the transformed additive and dominance matrices. It is not clear if this would make decryption easier.

(iii) Finally, the major principal components of the genotype dosage matrix are sometimes included as covariates, in place of or in addition to fitting a mixed model, in order to further control for population structure. That the *n* × *m* dosage matrix ***H*** has singular value decomposition ***H*** = ***UΣV***^***T***^, where ***U*** is the *n* × *n* orthogonal matrix of principal components, ***Σ*** is *n* × *n* diagonal and ***V***^***T***^ is *n* × *m* orthogonal. Thus ***F*** = ***PH*** = ***PUΣV***^***T***^. This means the principal components ***U*** of ***H*** are transformed to ***PU*** so that if necessary, the principal components of ***F*** may be calculated and included in the linear mixed model without explicitly including them as covariates to be transformed.

### Orthogonal Homomorphic Encryption

We propose that, if the orthogonal key ***P*** is appropriately sampled at random and independently of the untransformed data *D*(*I*) = {***y***, ***X***, ***H***}, then it homomorphically encrypts *D*(***I***) → *D*(***P***), sufficient to allow full mixed-model GWAS without revealing the plaintext.

The Pearson correlation between a standardised vector ***x*** and ***Px*** is

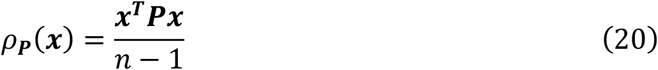

Thus, provided ***P*** is “far” from the identity matrix ***I*** then we expect *ρ*_***P***_(***x***) to be distributed like the correlation of two random vectors. An effective way to do this is to sample orthogonal matrices from the Steifel Manifold (i.e. the Haar measure over the orthogonal group)(Hoff 2009). This can be thought of as a uniform sampling distribution for orthogonal matrices (Anderson *et al.* 2005).

To investigate this experimentally, we sampled a 1000 × 1000 matrix *P*_1000_ using the R library “rstiefel”. This uses the following scheme to simulate an orthogonal *n* × *n* matrix (i) simulate an *n* × *n* matrix ***M*** whose entries are all iid *N*(0,1). (ii) compute the eigen-decomposition of the symmetric matrix ***M***^***T***^***M*** = ***Q***^***T***^***SQ*** where ***Q*** is *n* × *n* orthogonal and ***S*** is diagonal with positive entries. (iii) Return the orthogonal matrix ***P*** = ***MQ***^***T***^***S***^**−0.5**^***Q*** where ***S***^**−0.5**^ is the diagonal matrix whose elements are the reciprocals of the square roots of the eigenvalues.

Now the eigen-decomposition of an orthogonal matrix can be written as

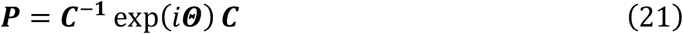

where ***C*** is a (non-orthogonal) matrix of eigenvectors and ***Θ*** is a diagonal matrix of angles, so that the eigenvalues exp(*i****Θ***) are pairs of conjugate complex numbers on the unit circle. Then, for *λ* real, define the set of orthogonal matrices ***P***(*λ*) = ***C***^**−1**^ exp(*iλ****Θ***) ***C***, which vary smoothly between ***P***(*λ* = 0) = ***I*** and ***P***(*λ* = 1) = ***P***.

Studying this set as *λ* varies lets us explore the encryption properties of a particular “linear direction” in the space of orthogonal matrices, starting at the identity matrix and passing through ***P***. [Incidentally, the set ***P***(*λ*) forms a subgroup of the orthogonal matrices, such that ***P***(*λ*)***P***(*μ*) = ***P***(*λ* + *μ*), with inverse ***P***(*λ*)^−1^ = ***P***(−*λ*). This subgroup is of course isomorphic to the real numbers under addition.]

The **Figure 3** shows the mean and standard deviation of the correlation *ρ*_***P***(***λ***)_(***x***) for a 1000 × 1000 matrix *P*_1000_ with 1,000 SNPs sampled from the CONVERGE study of major depressive disorder(Cai *et al.* 2015), for a subset of 1,000 randomly-sampled individuals. When *λ* = 0 then the correlations are all unity, as would be expected, but as *λ* increases we observe a damped oscillatory behaviour, with mean correlation of 0 at approximately *λ* = 1,2,3,…

**Figure 3.**
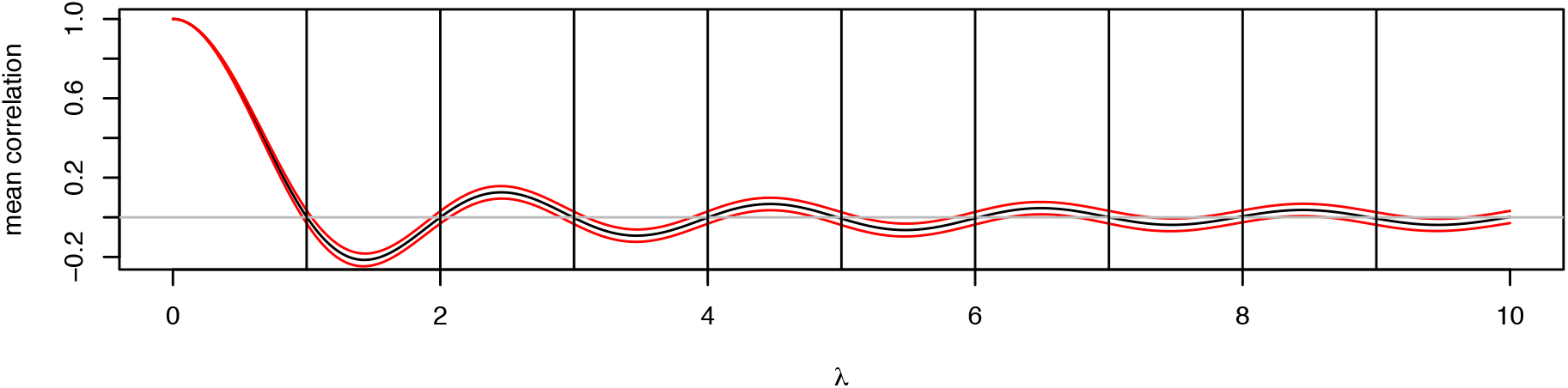
Correlation of unencrypted SNP dosages with encrypted versions as a function of λ. The black line shows the mean correlation ρ_P(λ)_(x) and the red lines the mean ± standard deviation, estimated from 1,000 individuals sampled from the CONVERGE study of major depressive disorder, at 1,000 randomly chosen SNP sites.

Thus, it is possible to sample a random orthogonal matrix such that on average there is no correlation between a random input vector of genotypes and its orthogonal transformation.

We applied these ideas to Human genotype dosages from the CONVERGE study of major depressive disorder in *n* = 10,465 individuals (Cai *et al.* 2015). We generated a random 10,465 × 10,465 orthogonal matrix ***P***_**10*k***_, which took about one hour with 2 cores and 8GB of RAM. Figure 4A shows the distribution of the correlations *ρ*_10*k*_(***x***) evaluated at 10,000 randomly chosen SNPs, after Z-transformation 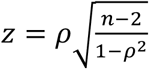, and **Figure 4B** shows the qqplot confirming the transformed correlations have a the expected null normal distribution. Figure 4C shows the distribution of standardised genotype dosages for a randomly selected SNP from that study, with values concentrated at the three modes corresponding to 0,1,2 reference alleles across the 10,465 individuals. Figure 4D shows the distribution of genotype dosages for the same SNP. It demonstrates that the values are close to normally distributed, centred at zero. Thus, transformed dosages are uncorrelated with their untransformed values, despite being a deterministic, invertible linear transformation of the latter. We return to this point later.

**Figure 4.**
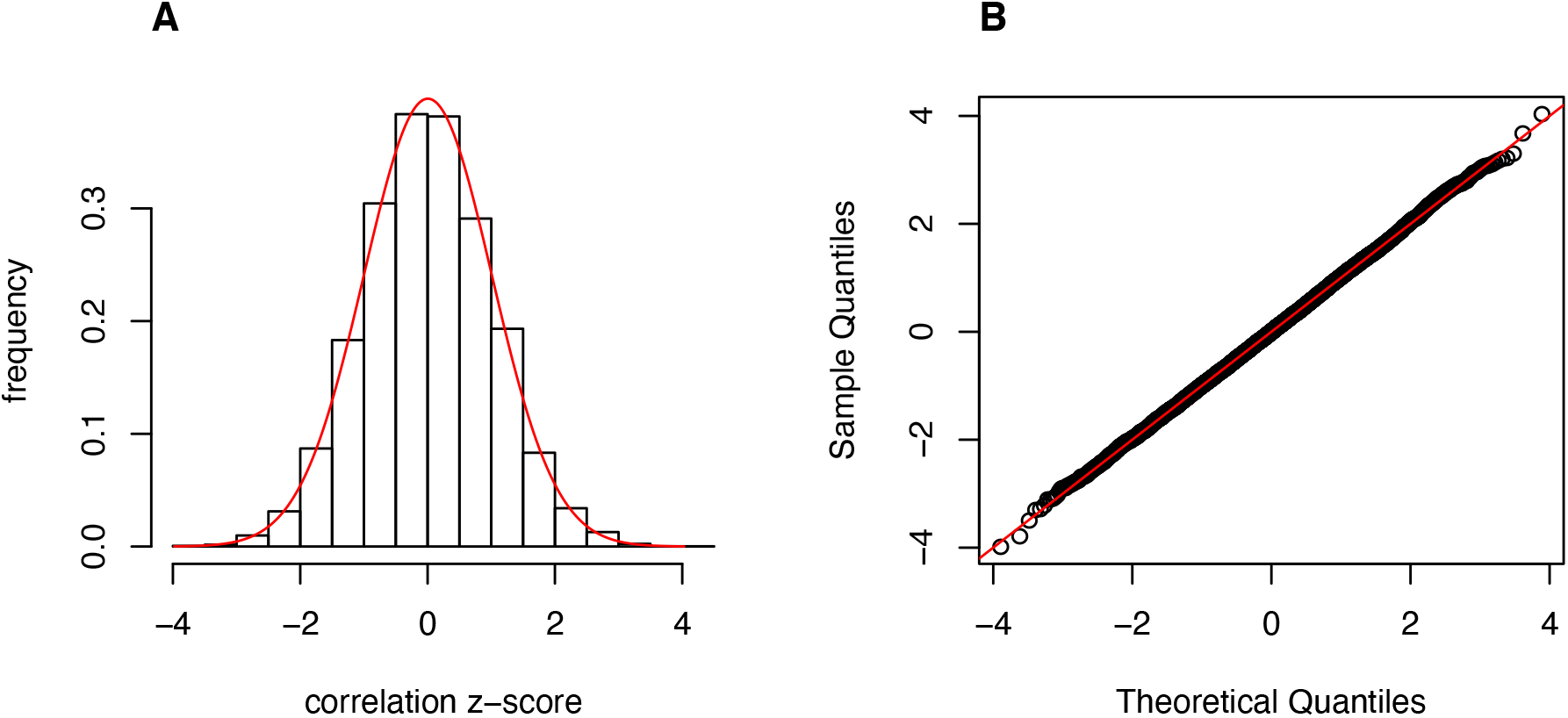
**A**: The distribution of Z-transformed correlations ρ_10k_(x) evaluated at 10,000 randomly chosen CONVERGE SNPs. The red line is the Normal density with the same mean and standard deviation. **B:** Normal quantile-quantile plot for the data in **A.**

A potential concern is that rounding errors might arise due to the very large dimension of the key ***P***. To test this, we computed ***P***^***T***^***P*** which should equal the identity matrix ***I***. When ***P*** = ***P***_**10*k***_The off-diagonal values (which should all equal 0) had typical magnitude 10^−11^, indicating the accuracy is acceptable. Nonetheless the average magnitude of off-diagonal elements drifts upwards as the dimension of the matrix increases – when ***P*** = ***P***_**1000**_ the magnitudes are typically only 10^−13^. Therefore, we might eventually encounter rounding issues when sampling very large orthogonal matrices, but not for matrix dimensions up to at least 10,000. One solution would be to divide the samples into randomly chosen blocks of 10,000 individuals, sample a different transformation matrix to encrypt each block, and then permute all transformed data so that the block structure is hidden.

Supplementary Material S1 contains R functions to generate random orthogonal matrices, encrypt genotype dosages and phenotypes, and to download and analyse an example publicly available mouse dataset from (Nicod *et al.* 2016) and perform a basic association study for platelet count on mouse chromosome 11 to demonstrate the methodology. These data and software are also available from UCL Figshare at https://rdr.ucl.ac.uk/account/home#/projects/76434.

### Applications of Orthogonal Genotype Encryption

It might be thought that orthogonal encryption is of little use, because both genotypes and phenotypes are transformed with the same orthogonal matrix, which must be known to those performing the transformation. However, there are uses for such a system. First, if the number of phenotypes is large (e.g. from a gene expression study) then it might be necessary to analyse the data on an insecure remote computing platform. Second, the encrypted data could be archived without special security concerns. Third, as we show next, it is possible to share and analyse federated independently-transformed data sets.

### Sharing Federated Transformed Genotype Data Sets

Suppose we wish to perform a federated mega-analysis on several genotype and phenotype sets. We assume that each set has first been imputed onto a common set of SNPs which are ordered consistently across data sets. Similarly, any covariates must be consistently defined and ordered across sets. Within each data set *D*_*t*_, with *n*_*t*_, subjects, an independent, private, orthogonal transformation is made using an *n*_*t*_ × *n*_*t*_ orthogonal matrix ***P***_***t***_ sampled at random to generate transformed data *D*_*t*_(***P***_***t***_)as above. We combine the shared transformed data by stacking them top of each other. Thus, for three sets we have:

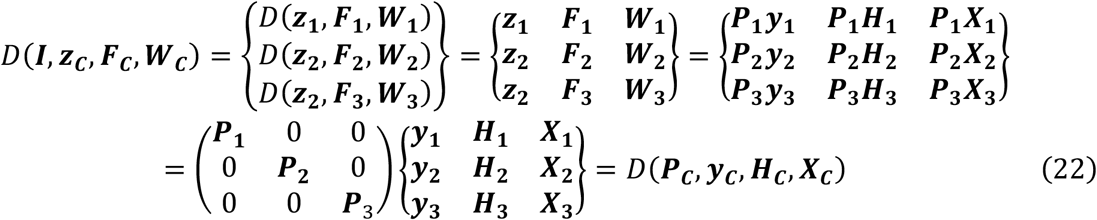

where the subscript ***C*** denotes the combined data and where the individual orthogonal matrices have been combined in a block-diagonal manner:

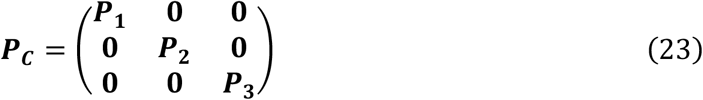

***P***_***C***_ is orthogonal ∑_*t*_ *n*_*t*_ × ∑_*t*_ *n*_*t*_ and hence the combined data can be analysed *as if* it were a single untransformed data set that had been encrypted using ***P***_***C***_. However, in reality each laboratory contributing a dataset *D*_*t*_ independently encrypts their data using their private key ***P***_***t***_ before sharing it.

Similarly, a dataset could also be subdivided into subsets (e.g. into male vs female subjects) and each part encrypted separately so that sub-analyses could be performed, and the subsets distributed separately. We emphasise that for federated analysis to work, it is necessary for the parties to agree in advance on a common set of SNPs and covariates.

### Removing Duplicates and Close Relatives: Dual Encryption

One potential difficulty when sharing encrypted data is the possibility of duplicates or close relatives occurring in different cohorts. Because HEGP disguises genetic relationships it would not be possible to identify duplicates in the shared ciphertexts. Whilst there are simple practical ways of eliminating individuals with identical IDs in different studies (eg first sharing the hashes of their sample IDs) or with identical genotypes at a small set of test SNPs (by sharing hashes of their genotype vectors), these methods would fail if the IDs were different or if the test genotypes differed even slightly (as might happen if the same samples were genotyped twice).

A solution would be for all parties to first agree on a restricted subset of *N*_*R*_ common test SNPs (say no more than 100 common SNPs chosen genome-wide at random). Each party computes their normalised plaintext ***H***_***R***_ restricted to just these SNPs, and they share the dual ciphertext ***F***_***R***_ = ***H***_***R***_***P***_***R***_, where ***P***_***R***_ is a random *N*_*R*_ × *N*_*R*_ orthogonal key, instead of sharing ***F*** = ***PH***.

Importantly, ***F***_***R***_ defines a dual form of encryption that has complementary properties to those of ***F***; for the dual GRM

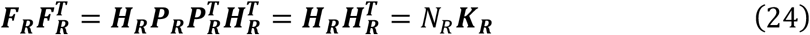

is the same as the plaintext GRM and instead the SNP correlation matrix is scrambled: dual encryption is therefore useless for genetic association. Close relatives and duplicates may then be identified from the GRM ***K***_***R***_, and then agreement reached on a revised subset of individuals from each study to be shared using the original scheme of encryption applied to all SNPs. However, it should be pointed out that sharing information in any way that reveals relationships between people is inherently risky.

### How Secure is Orthogonal Encryption?

Can we determine ***P*** given only *D*(***P***)?. Although we have shown that ***PH*** is uncorrelated with ***H***, we have not shown this renders the transformation truly secure. Since the encryption and decryption keys are essentially the same, this form of encryption has very different properties from public-key methods. Orthogonal encryption is certainly insecure for certain choices of ***P***. As **Figure 3** shows, any orthogonal matrix close to the identity matrix is clearly a poor choice, so one should restrict attention to random orthogonal matrices sampled from either the Steifel Manifold or using another scheme with similar sampling properties. One should also check that the mean of the correlations of the columns of ***F*** with the columns of ***H*** is close to zero.

It is obvious that any permutation of the rows of any key will transform the phenotype and genotypes in the same way, and so are functionally equivalent. Consequently, orthogonal permutation matrices are useless as keys. However, it also means that any permutation of any good key is also a good key.

The singular value decomposition of the unencrypted *n* × *m* dosage matrix ***H*** has ***H*** = ***UΣV***^***T***^, where ***U*** is *n* × *n* orthogonal, ***Σ*** is *n* × *n* diagonal and ***V***^***T***^ is *n* × *m* orthogonal. Thus ***F*** = ***PH*** = ***PUΣV***^***T***^, so the rotation ***U*** is simply replaced by another rotation ***PU***. If ***P*** is truly random then we seek ***U*** given ***PU*** which appears to be hard problem, since ***PU*** resembles another random orthogonal transformation.

Next we discuss strategies that might be used to decrypt the data, in likely increasing order of effectiveness:

#### Brute Force

(a) We first tried sampling random decryption keys using the rustiefel method. Each key contains *n*(*n* + 1)/2 independent double precision numbers, each of which can take about 10^20^ possible values. We defined a distance function between matrices equal to the mean of the absolute difference between each pair of corresponding elements (i.e. the L1 norm), to compare the plaintext genotype matrix to an attempted decryption. We defined a “good” key as one that gives a mean distance of less than 0.4 between the genotype matrix and the attempted decryption. Empirically this upper limit gave results that are visually fairly close to the original, at least for small datasets. Extrapolating from small matrices, we estimated a lower bound on the number of attempts required for solving an *n* × *n* key, of one “good” key generated per 10^*n*−1^ incorrect keys. thus, if *n* = 100, at least 10^99^ keys have to be tried before a good one is found. Interestingly, even for an 8 × 8 matrix we could not a key that regenerated the plaintext, and even ‘good’ keys do not reflect the underlying genotypes fully.

Generating orthogonal random keys is computationally expensive – the computational complexity of the Stiefel manifold is *O*(*n*^3^); if *n* = 100 a few hundred keys can be generated and evaluated per second on one CPU core. Our estimated bound suggests that it would take in the order of 10^92^ CPU hours to get close to a solution. Larger keys of realistic size take significantly longer – e.g when *n* = 10,000, a single key takes about one CPU hour to generate. Rather than generating orthogonal keys, a naive brute force attack where potential keys are randomly selected would be even slower because the search space becomes much larger including all non-orthogonal matrices. Thus our method takes a great deal of CPU power to guess large orthogonal matrices.

These experiments show that there is no consistent relationship between generated keys and their decryption outcomes using this simple metric. Moreover, as the distance function we used is defined in terms of distance to the known plaintext, it only works when we know the end result. In reality, an attacker would have to use a less accurate score function. The number of possible permutations of the result matrix is so large that it is not feasible to brute force an attack without a method optimized to compute orthogonal matrices while optimizing for a metric that has an open-ended end result.

#### Exploiting non-Gaussian Distributions of Genotype Dosages

Another potential attack, that exploits specific features of the problem, is as follows. We note that the SNP identities (genomic positions) need to be distributed with the data in order to interpret the biology of any GWAS hits. Population allele frequencies for SNPs are generally available, and so for a SNP *j* with frequency *π*_*j*_ that is in Hardy-Weinberg equilibrium, we expect to observe genotype dosages in the proportions

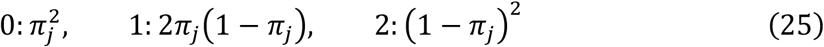

After standardisation the dosages will be rescaled but will still be tri-modal with modes *d*_*j*0_, *d*_*j*1_, *d*_*j*2_ that are completely determined by *π*_*j*_ and the constraints that the standardised dosages have mean=0, variance=1 and that *d*_*j*0_ − *d*_*j*1_ = *d*_*j*1_ − *d*_*j*2_.

Consequently, we might seek an orthogonal matrix approximation ***Φ*** ≈ ***P***^**−1**^ = ***P***^***T***^ that maps ***F*** → ***ΦF*** with columns such that each has the frequency distribution close to that predicted by HWE. That is, for each SNP, the decrypted genotype dosages will be expected to have been sampled from a distribution with modes at *d*_*j*0_, *d*_*j*1_, *d*_*j*2_ corresponding to the rescaled dosages 0,1,2 (like **Figure 1C**), which can be modelled using a kernel density estimate

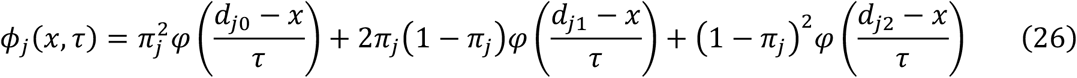

where *φ*(*z*) is a standard normal density kernel and *τ* is the standard deviation of the kernel. Then we seek an orthogonal matrix *Φ*\* that maximises

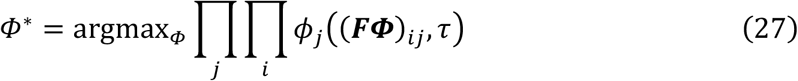

We also require *τ* to be small in order to concentrate the data around the modes. However, if the plaintext dosages were imputed then they might well not be exactly integral, so it is necessary that *τ* > 0 but still as small as reasonably possible.

Equation (27) describes a non-convex and non-linear objective function. One potential approach to minimisation is via robust non-convex optimisation based on the Cayley transform(Bertsimas *et al.* 2010; Wen and Yin 2013). Whether such an attack is feasible is unclear: the space of *n* × *n* orthogonal matrices has dimension *n* (*n* − 1)/2, so if *n* = 10^4^ the minimisation is over 4.995 × 10^7^ ≈ 50 million free parameters. There are likely to be local minima. It is also unclear if the minimiser *Φ*\* is unique, or that the true answer necessarily minimises this quantity (by unique we mean if two distinct solutions *Φ*\*\*, *Φ*\* exist then they are permutations of each other.

Fast Independent Components Analysis (FastICA) (Hyvärinen and Oja 1997), is another method that attempts to split non-Gaussian signal from Gaussian noise. FastICA(Hyvärinen and Oja 1997)(Hyvärinen and Oja 1997)(Hyvärinen and Oja 1997)(Hyvärinen and Oja 1997)(Hyvärinen and Oja 1997) finds an orthogonal transformation to map the data onto “interesting directions” such that the projections of the data are strongly non-Gaussian along these directions: in our case, we seek directions in which the distributions are trimodal. In this context, FastICA may be thought of as maximising a different function from the likelihood with a particular choice of optimisation algorithm. However, we found that applying the implementation in the “fastICA” R package does not improve on our random brute force attacks. We configured FastICA to produce an orthogonal matrix of the same size as the encryption key and computed the distance score of the resulting matrix. We found these scores were much higher than the best keys generated during the brute-force attack. This is the case whether using a random initial matrix, or providing an already generated key with a relatively good score. **Table 1** shows the results of applying FastICA to the seven best keys from the brute-force attempt on a 4×4 key, with a 4×636 dataset. The per-entry error in the decrypted data of 0.079 on average, which is quite good, while anything greater than 0.4 is unusable.

**Table 1.**
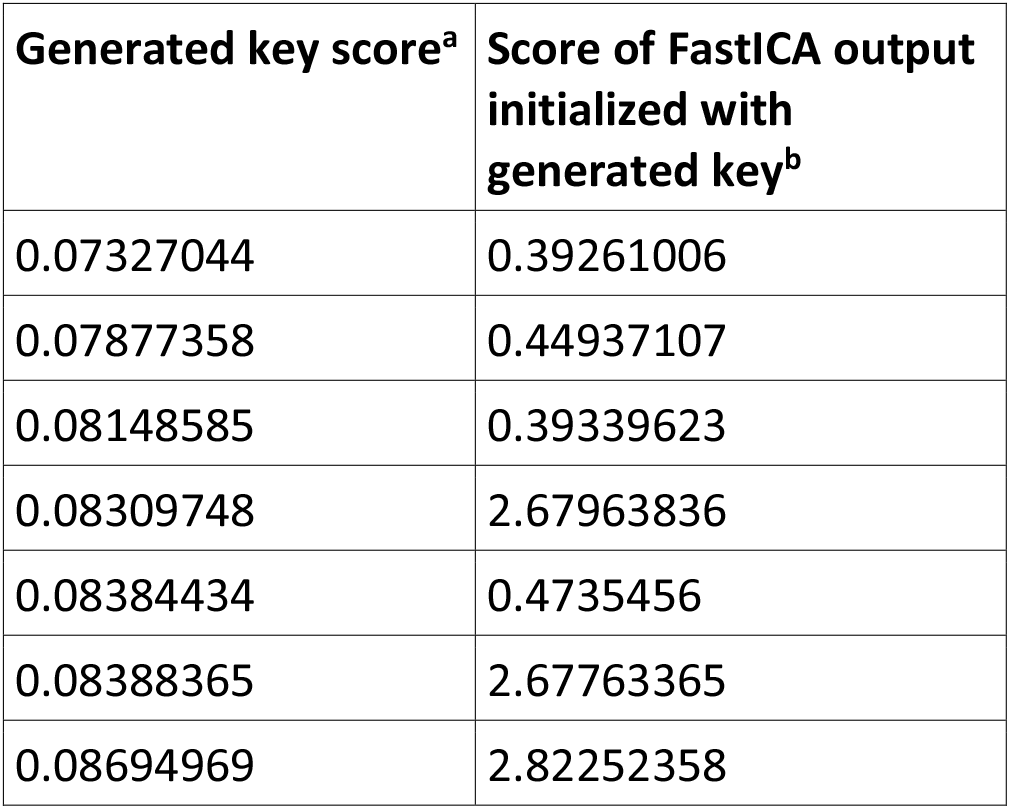
Examples of attempted decryption using FastICA. Seven 4×636 genotype matrices were first encrypted and then attempts at decryption made either by (a) brute force random attempts or (b) FastICA. The score is the L1 distance between matrices

| Generated key score<sup>a</sup> | Score of FastICA output initialized with generated key<sup>b</sup> |
| --- | --- |
| 0.07327044 | 0.39261006 |
| 0.07877358 | 0.44937107 |
| 0.08148585 | 0.39339623 |
| 0.08309748 | 2.67963836 |
| 0.08384434 | 0.4735456 |
| 0.08388365 | 2.67763365 |
| 0.08694969 | 2.82252358 |

As FastICA attempts to maximize non-Gaussianity in the decrypted data, these results imply that non-Gaussianity does not describe the desired decrypted data sufficiently uniquely. While the decrypted data is non-Gaussian, there are many other transformations the encrypted data that also produce highly non-Gaussian results.

Another form of mathematical optimization is constrained convex programming, where constraints could be imposed to ensure the decrypted genotypes take plausible values. The main difficulty with applying convex programming (and linear programming, which also handles constraints) is the choice of a suitable objective function to minimise. There is good reason to believe convex programming cannot produce good results. Optimizing a key to improve its decryption results would entail finding a path through the *n*-dimensional space of rotations, choosing both a correct direction to rotate in, and degree of rotation at each step. Specifically, the score function is not locally convex, and any naive optimization attempt is bound to fall into local minima. Similarly, gradient descent is also unlikely to be useful, as each iteration would require calculating a number (linear to the size of the key) of matrix multiplications (of the entire dataset with the key at each step).

#### Compression

A restatement is that the plaintext is highly compressible (at least, if all the genotypes are integral), so we might instead seek

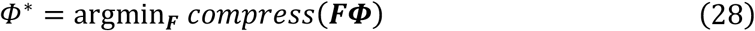

where “compress” is some program like gzip. Again, we do not know if the most compressible encoding of the genotypes is identical to the true answer, or whether this could be computed efficiently. We expect it would be very slow for large datasets.

#### Pedigrees

If all the individuals in the study are from a set of known pedigrees (for example a large set of trios) then the expected plaintext GRM 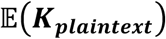 is known (ie entries for full sibs and parent-offspring will be ½, unrelateds will be 0, etc) and we can assume the samples are ordered so that the matrix is block-diagonal. Then the original and encrypted GRMs are linked via the approximation

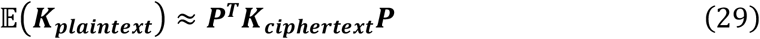

which is a system of *n*(*n* + 1)/2 quadratic equations in *n*(*n* + 1)/2 independent unknowns (the number of degrees of freedom in an *n* × *n* orthogonal or symmetric matrix). Thus an approximation to ***P*** could be obtained and might be a useful initial guess for further refinement, if *n* is small. When *n* is large the problem has exponential time complexity (Grigoriev and Pasechnik 2005). Moreover, any permutation of the ordering of pedigrees that left ***K***_***plaintext***_ unchanged would have the same solution so it would be impossible to assign phenotypes to pedigrees uniquely. Finally, if everyone is unrelated then 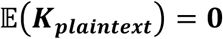 and the method would not work.

#### Incremental Decryption

Another way of thinking about the effects of the group of orthogonal transformations is that they define sets of equivalence classes of data sets *D*. That is, two sets *D*_1_, *D*_2_ are equivalent if there exists and orthogonal matrix ***P*** such that *D*_1_(***P***) = *D*_2_ i.e. that maps one to the other. The transitive property of the group of orthogonal matrices means that there is always an orthogonal matrix that will transform any pair of datasets provided they are in the same equivalence class. All data sets in the same equivalence class have the same likelihood so these classes can be thought of as likelihood contours in a high-dimensional space.

This suggests another attack on the problem: find a series of *N* incremental orthogonal transformations that successively resolve individuals by “factorising along the contour”. That is, we seek a sequence of orthogonal keys {***Φ***_***k***_} and partially decrypted genotype matrices {*F*_*k*_} such that (a) ∏_*k*_ ***Φ***_***k***_ = ***P***^***T***^, (b) ***F***_***k***_***Φ***_***k***_ → ***F***_***k*+1**_ with ***F***_**1**_ = ***F***,***F***_***N***_ = ***H***. There certainly exist infinitely many orthogonal keys that decrypt any subset of individuals. Suppose we want to decrypt the first *k* individuals. Then if ***Q***_***n***−***k***_ is any *n* − *k* × *n* − *k* orthogonal matrix and ***I***_***k***_ is the *k* × *k* identity, and we partition ***P*** = [***P***_***k***_|***P***_***n***−***k***_], such that ***P***_***k***_ is the *n* × *k* matrix comprising the first *k* columns of ***P***, and ***P***_***n***−***k***_ is the last *n* − *k* columns, then

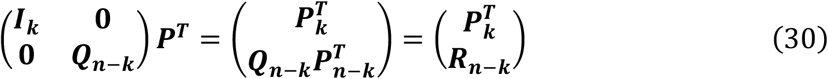

where ***R***_***n***−***k***_ is any (*n* − *k*) × *n* orthogonal matrix, will decrypt just the first *k* individuals. Thus, a sequence of matrices of the above form would decrypt the data. Using this scheme, in principle one could either try to decrypt individuals one-by-one in *N* = *n* steps or use a divide-and-conquer strategy with *N* ≈ log_2_ *n* more difficult steps. Of course, since we do not know ***P***^***T***^ this merely proves existence: it is not clear that this type of approach is intrinsically better than trying to estimate ***P***^***T***^directly – one still needs to estimate each column of ***P***^***T***^, a non-trivial task.

#### Private Variants

There is one clear-cut weakness to orthogonal encryption, which occurs when ultra-rare private variants are present. Suppose a SNP *j* is private to the individual *i*. Then the genotype dosages for this SNP (column *j* of ***G***) comprises *n* − 1 zeros and one non-zero value, say 1 at row *i*. After standardisation this pattern is preserved in ***H*** although the numerical values are now scaled so the *j*’th column mean is zero and its variance is unity. The column *j* of ***PH*** is then

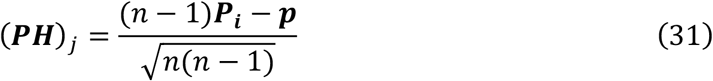

i.e. a linear combination of column *i* of ***P*** and a fixed vector ***p*** equal to the row sums of ***P***. This reveals the decryption key for individual *i* if ***p*** can be guessed.

Thus, in an extreme case, should every individual carry a private variant, or equivalently if *n* covariates were defined that uniquely identify each individual, then the system can be attacked successfully. While this is an unlikely situation in practice, and one that could easily be avoided, it does suggest that an attack focussed on lower frequency variants might be able to extract useful information. Further, once an individual has been decrypted in this way then close relatives might be more easily identifiable as well.

Equation (30) shows how private variants could be factored out, leaving a smaller orthogonal key representing common variants still to be discovered. Note however that factoring out those individuals with private variants does not reveal useful information about other unrelated individuals because the remaining columns of ***P*** are in the subspace orthogonal to that spanned by the factored columns. In addition, population allele frequencies need not perfectly match those in the sample, so it is not necessarily clear which variants are in fact private. Moreover, there is no relationship between allele frequency and the correlation between cyphertext and plaintext dosages, as is shown in **Figure 5,** which plots the squared correlations as a function of allele frequency for simulations.

**Figure 5.**
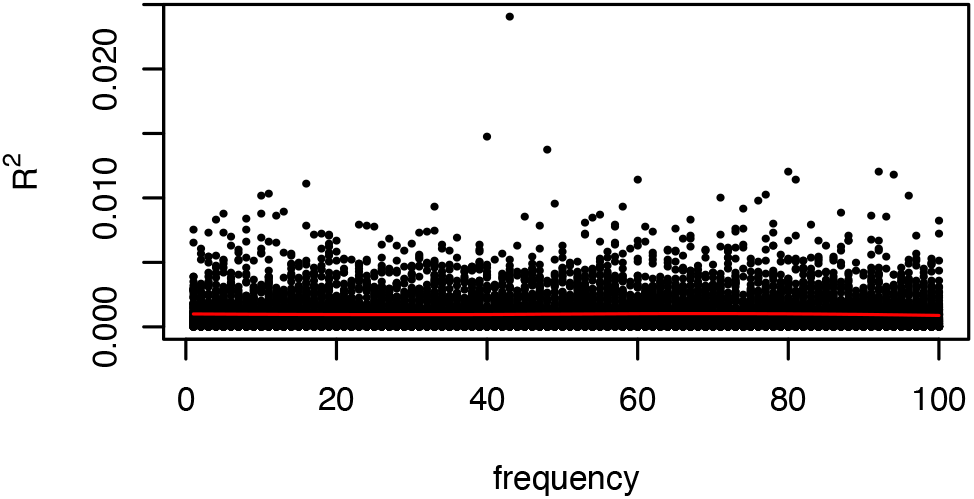
Correlation R^2^ of plaintext and ciphertext dosages as a function of minor allele frequency. Simulations are of genotypes for 1000 subjects with minor allele frequencies in the range (1…100)/1000. Each black dot represents one vector of genotypes. y-axis: squared correlation R^2^, x-axis: allele frequency. Red line is the smoothed moving average of R^2^.

### Practical Implementation Issues

We tested our encryption scheme on 10,640 individuals from the CONVERGE study of major depressive disorder (Cai *et al.* 2017), and on the smaller mouse dataset of 1,329 individuals and 19,877 SNPs from (Nicod *et al.* 2016) for platelet counts on mouse chromosome 11, that are publicly available as described in Supplementary Data 1. We use the mouse data for the majority of the analyses in this study so that users may replicate our analyses by downloading the data and code.

HEGP leaves the calculation of genetic association unchanged, so should analyse ciphertext in the same execution time as with plaintext. Software that runs off genotype dosage data should run altered since the rotated data are dosage-like. HEGP cannot deal with missing data, and which should be imputed first. Another restriction is that it is impossible to analyse subsets of the individuals (eg all those of one sex) once they have been encrypted, unless each subset was encrypted separately. However, if a covariate specifying sex is also encrypted then it would be possible totake sex into account when fitting the model.

The simulation of very large orthogonal keys (e.g. for hundreds of thousands of individuals) might also present technical difficulties. A simple solution would be to first permute the rows of *D*, ad then group them into a maximum of about 1,000 − 10,000 individuals per group, sample an independent orthogonal key to encrypt each group separately, as described above. The initial permutation would enhance the security of the data by separating potentially similar individuals. [Permutations are also orthogonal transformations, although in isolation they are useless encryptors as they rearrange phenotype and genotype identically.]

For the human data, we encrypted the phenotype and genotype dosages in 10 groups of 1000 individuals plus a final block of 664. We computed association across 160k SNPs using both unencrypted and encrypted dosages. The correlation between the logP values of the tests of association was 0.999. The average absolute difference between the logP values was 0.002. All calculations were performed in R using standard matrix arithmetic. Bearing in mind that usually only the first two decimal places of a logP value are of interest when interpreting the significance of genetic association, we conclude the numerical inaccuracies introduced by the encryption are negligible.

For the mouse data the mean absolute difference in logP values for simple association was 6.406e-03, with a maximum of 3.775e-02. We also implemented the mixed model (Equation 13) to confirm that heritability estimates and association p-values are numerically stable after encryption. For the mixed model the mean absolute difference was 3.141e-03, maximum 2.635e-02. The mixed-model heritability estimated from the unencrypted data was 0.02534315, compared to 0.025049 after encryption, a discrepancy of 1.1%. We conclude that HEGP does not noticeably affect GWAS results.

### Quantile Normalisation to Improve Security

**Figure 1D** shows the distribution of ciphertext dosages for a given SNP is almost Gaussian. This suggests quantile normalising the ciphertext might improve security. In this scheme, the values in each column of ***F*** are first ranked and then replaced by their corresponding standard Normal quantiles. After quantile normalisation the the columns of ***F*** contain different permutations of the Normal quantiles of 1/(*n* + 1)…*n*/(*n* + 1) that respect the rank orders of the original ciphertext for each column, applying a small non-linear perturbation to the encrypted genotypes, ***F*** → ***F***_***q***_. Attacks that exploit non-normality in the encrypted data would be frustrated, potentially increasing security. A further refinement might iterate an alternating sequence of independent rotations and quantile normalisations.

We evaluated the effects of quantile normalisation on the ciphertext mouse genotypes and platelet phenotypes. First, the mean absolute discrepancy for mixed-model association logP values for the plaintext *vs* HEGP ciphertext was 0.003141, (maximum 0.0263), and the overall correlation of logP values was 0.999: a close agreement. The mean absolute difference between the plaintext and ciphertext dosages (i.e. L1 norm) ⟦***H*** − ***P***^***T***^***F***⟧ was 3.561 × 10^−9^, maximum 1.773 × 10^−6^. Thus HEGP alone induces only negligible reductions in accuracy of association statistics and genotypes. However, after encryption and quantile normalisation, the mean logP discrepancy rose slightly to 2.402 × 10^−1^, maximum 2.257 × 10^−1^, but the correlation was still over 0.99. Similarly, the estimated heritability changed 1.3% from 0.02472 to 0.0250. However, the mean absolute error in the decrypted quantile-normalised standardised genotype dosages ⟦***H*** − ***P***^***T***^***F***_***q***_⟧ rose to 0.03585 (i.e. mean discrepancy 18%), maximum 0.06980.

Our interpretation of this observation is that plaintext dosages correspond to a very special choice of coordinates where the standardised genotype dosages for a SNP are concentrated on three modes depending on the SNP allele frequency. Any random rotation of the genotypes produces coordinates such that the ciphertext dosages closely resemble a Gaussian sample. After rotating into such a coordinate frame it is then possible to make small non-linear perturbations that have little effect on association statistics or heritability but degrade the decryption back into the true coordinate system.

We also explored adding further security by quantile normalising and rounding the encrypted dosages. As would be expected, there is a trade-off between the number of significant digits retained after rounding and the accuracy of association and decryption.

### Logistic Regression

So far we have considered quantitative traits with Normally distributed errors, analysed in a mixed model framework. Whilst case-control studies (ie where the phenotype *y* ∈ {0,1}) are often analysed as if they were quantitative traits, under some circumstances it is preferable to use logistic regression, where

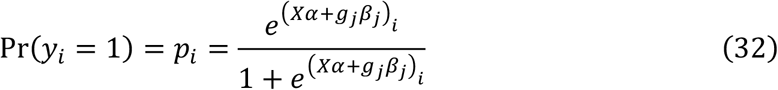

where (***Xα*** + *g*_*j*_*β*_*j*_) is the *i*th element of the vector ***Xα*** + *g*_*j*_*β*_*j*_. Write ***X***_***j***_ = [***X***|***g***_***j***_] and ***α***_***j***_ = [***α***|*β*_*j*]_. The likelihood for the data at SNP *j* is

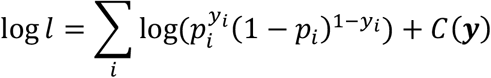

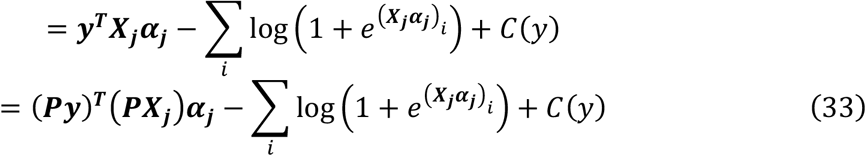

for any orthogonal matrix ***P***, and where *C*(***y***) is a function of ***y*** only that can be ignored when maximising the likelihood. Thus, the likelihood function comprises two components, namely ***y***^***T***^***X***_***j***_***α***_***j***_, which is invariant under orthogonal transformation, and 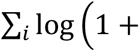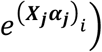, which is not invariant, instead transforming like 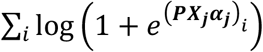. However, only the first component involves both the dependent and independent variables. This component is shared with the log-likelihood for the Normal linear model, which is why fitting a linear model to case-control data generates p-values resembling those from logistic regression. It should be clear that case-control data (i.e. *y*_*i*_ ∈ {0,1}) is no longer of the same form after an orthogonal transformation, so strictly speaking the likelihood no longer represents a logistic model after transformation. Nonetheless we can attempt to estimate parameters by maximising the transformed likelihood (Equation 33).

We fitted the logistic log-likelihood model to simulated SNP data, using untransformed and orthogonally transformed data in order to assess the change in maximum likelihood parameter estimates under transformation. We found that the estimates changed considerably and therefore orthogonal encryption is not homomorphic for logistic regression, for which we therefore recommend methods such as (Wang *et al.* 2015).

### The Mixed-Model Linear Transformation as an alternative Encryptor

Are any non-orthogonal transformations suitable for homomorphic encryption? The mixed-model transformation *A*^−1^ shares some, but not all, of the invariant properties of the orthogonal group. If we set

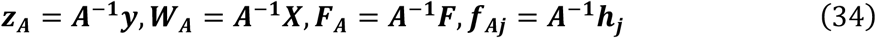

Then the Var(***z***_***A***_) = ***I*** and the log-likelihood transforms thus:

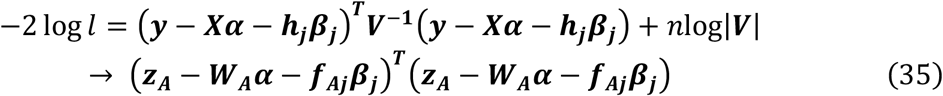

Thus the log-likelihood is preserved so we can extract the mixed-model GWAS p-values as before. Moreover, ***A***^**−1**^ has *n*^2^ free parameters, compared to *n* (*n* − 1)/2 for ***P***, so the decryption problem is presumably harder. Furthermore, it is easily seen that ***A***^**−1**^ may be replaced by ***PA***^**−1**^ for any orthogonal ***P*** making the decryption harder still. However, there is some loss of information: It is no longer possible to estimate the variance components 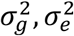 nor the heritability *h*^2^. Furthermore, a federated analysis along the lines described above would not give exactly the same p-values as would orthogonal transformation followed by a mixed-model transformation applied to the combined dataset, because each component study has been transformed separately without guaranteeing the federated transformed GRM is also the identity; the structure of the federated variance matrix will be of the form

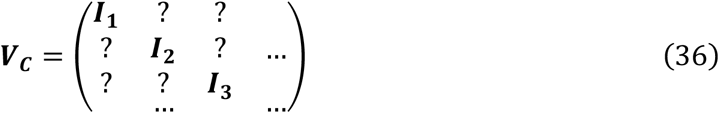

Lastly, linkage disequilibrium between the SNPs is no longer conserved:

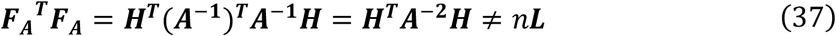

(using the fact that ***A***^**−1**^ is symmetric).

## Discussion: A Community Challenge

HEGP has many desirable properties for quantitative genetics. It preserves linkage disequilibrium between genetic variants, and key association statistics including heritability between variants and phenotypes, while obscuring relationships between individuals. However, we do not yet fully understand when HEGP is cryptographically secure. Where private variants are available, decryption is straightforward. While it is simple to remove low frequency variants and therefore protect against this weakness, the larger question of security remains. We have sketched out several potential attacks but so our investigations have not found a workable method. To settle this question, one would need either to find an efficient inversion algorithm - perhaps a version non-convex minimisation under constraints(Bertsimas *et al.* 2010) - that recovers the correct genotypes accurately, or alternatively to show there are too many incorrect “genotype-like” decoy solutions far from the true answer, and that therefore the problem is essentially non-invertible. It is likely that the inversion problem might be solvable for small data sets, but much harder for larger ones.

Orthogonal encryption also has the potential weakness that the key space is continuous; in conventional crypto, a small change in the key used leads to a completely different ciphertext. In contrast a small change to an orthogonal key leads to small changes in the ciphertext. However, at this point, we know of no algorithm that can exploit this. We found that transformed genotypes closely resemble samples from a Normal distribution, and so can be replaced by exact Normal quantiles with only small effects on accuracy. Hence we can certainly protect the ciphertext from attacks that rely on Non-Gaussianity.

The hardness of the inversion problem depends not only on avoiding private variants, but on choosing a good key. Those sampled from the Steifel manifold work well at obscuring correlations between plaintext and ciphertext genotypes, such that - as measured by mean correlation across all sites - transformed individuals do not resemble the originals more closely than do simulated individuals with matched allele frequencies. However, it is possible that other measures of genetic similarity between individuals might not be randomised to the same extent.

Thus more work is needed to determine precisely when random orthogonal keys are cryptographically secure. **We submit this problem as an open challenge to the community.**

While HEGP lacks mathematical proof of security compared to normal crypto schemes, most schemes are broken due to weaknesses in implementation (bad random number generators, sidechannel attacks, etc.), not algorithm. HEGP has the advantage of an extremely simple algorithm, and is probably immune to sidechannel attacks (and to an extent social engineering and rubber-hose cryptanalysis).

Given our current knowledge, we claim that random orthogonal keys provide an encryption scheme where it is - at the least - very difficult to recover individual genetic or phenotypic data. This is at least equal to the level of security of a date shift of medical records which is also not completely secure but makes it difficult for researchers to identify an individual if they do not intend to do so.

Thus, should an effective attack be discovered, orthogonal keys still offer “pretty good genetic privacy” in the sense that they would prevent straightforward copying of information about individuals’ genotypes. We argue that routine orthogonal transformation of genotypes and phenotypes, in combination with existing legal protocols, would enhance security, increase collaboration and data sharing, and thereby accelerate progress.

In summary, we have shown how to make a distinction between public information about genetic architecture and allelic effects, and private information about individuals. This general principle could be applied more widely. We mention two examples:

First, to the extent that medical records can be analysed in a linear modelling framework with a suitable design matrix, orthogonal encryption offers a means to perform federated analyses on orthogonally encrypted medical records.

Second, genetic improvement of crops and farm animals could be accelerated. Whilst some germplasm and genetic variation data are in the public domain, commercial breeders are developing new varieties and breeds and have extensive proprietary genetic and phenotypic data that could be usefully shared using HEGP, so that alleles conferring a beneficial trait could be discovered and published without revealing the genomes of proprietary germplasm under development.

Such a move - towards the idea that an allele’s effects are public property whilst an individual’s genotypes are private - is more important than the encryption mechanism used to attain it.

## Acknowledgments

We thank Rob Williams, the *Genetics* Editors and the anonymous reviewers for valuable comments.

## References

Anderson T. W., Olkin I., Underhill L. G., 2005 Generation of Random Orthogonal Matrices. SIAM J. Sci. Stat. Comput. 8: 625–629.

Azencott C. A., 2018 Machine learning and genomics: Precision medicine versus patient privacy. Philos. Trans. R. Soc. A Math. Phys. Eng. Sci. 376: 20170350.

Bertsimas D., Nohadani O., Teo K. M., 2010 Nonconvex robust optimization for problems with constraints. INFORMS J. Comput. 22: 44–58.

Bonte C., Makri E., Ardeshirdavani A., Simm J., Moreau Y., et al., 2018 Towards practical privacy-preserving genome-wide association study. BMC Bioinformatics 19: 537.

Cai N. N., Bigdeli T. B. T. B., Kretzschmar W., Lei Y., Liang J., et al., 2015 Sparse whole-genome sequencing identifies two loci for major depressive disorder. Nature 523: 588–591.

Cai N., Bigdeli T. B., Kretzschmar W. W., Li Y., Liang J., et al., 2017 11,670 whole-genome sequences representative of the Han Chinese population from the CONVERGE project. Sci. data 4: 170011.

Cho H., Wu D. J., Berger B., 2018 Secure genome-wide association analysis using multiparty computation. Nat. Biotechnol. 36: 547–551.

Grigoriev D., Pasechnik D. V., 2005 Polynomial-time computing over quadratic maps i: Sampling in real algebraic sets. Comput. Complex. 14: 20–52.

Hansson M. G., Lochmüller H., Riess O., Schaefer F., Orth M., et al., 2016 The risk of re-identification versus the need to identify individuals in rare disease research. Eur. J. Hum. Genet. 24: 1553–1558.

Hoff P. D., 2009 Simulation of the matrix Bingham-von Mises-Fisher distribution, with applications to multivariate and relational data. J. Comput. Graph. Stat. 18: 438–456.

Hripcsak G., Mirhaji P., Low A. F. H., Malin B. A., 2016 Preserving temporal relations in clinical data while maintaining privacy. J. Am. Med. Informatics Assoc. 23: 1040–1045.

Hyvärinen A., Oja E., 1997 A Fast Fixed-Point Algorithm for Independent Component Analysis 1 Introduction. Most 9: 1483–1492.

Jagadeesh K. A., Wu D. J., Birgmeier J. A., Boneh D., Bejerano G., 2017 Deriving genomic diagnoses without revealing patient genomes. Science (80-.). 357: 692–695.

Kang H. M., Zaitlen N. A., Wade C. M., Kirby A., Heckerman D., et al., 2008 Efficient control of population structure in model organism association mapping. Genetics 178: 1709–1723.

Nicod J., Davies R. W. R. W., Cai N. N., Hassett C., Goodstadt L., et al., 2016 Genome-wide association of multiple complex traits in outbred mice by ultra-low-coverage sequencing. Nat. Genet. 48: 912–918.

Pasaniuc B., Price A. L., 2017 Dissecting the genetics of complex traits using summary association statistics. Nat. Rev. Genet. 8: 117–127.

Sim J. J., Chan F. M., Chen S., Tan B. H. M., Aung K. M. M., 2019 Achieving GWAS with Homomorphic Encryption. ArXiV.

Tkachenko O., Weinert C., Schneider T., Hamacher K., 2018 Large-Scale Privacy-Preserving Statistical Computations for Distributed Genome-Wide Association Studies. In: Proceedings of the 2018 on Asia Conference on Computer and Communications Security,, pp. 221–235.

Wang S., Zhang Y., Dai W., Lauter K., Kim M., et al., 2015 HEALER: Homomorphic computation of ExAct Logistic rEgRession for secure rare disease variants analysis in GWAS. Bioinformatics 32: 211–218.

Wen Z., Yin W., 2013 A feasible method for optimization with orthogonality constraints. Math. Program.

Yang J., Lee S. H., Goddard M. E., Visscher P. M., 2011 GCTA: a tool for genome-wide complex trait analysis. Am. J. Hum. Genet. 88: 76–82.

